# N^α^-acetylation of EsxA is required for mycobacterial cytosolic translocation and virulence

**DOI:** 10.1101/2020.01.08.899369

**Authors:** Javier Aguilera, Chitra B. Karki, Lin Li, Salvador Vazquez-Reyes, Qi Zhang, Chenoa D. Arico, Hugues Ouellet, Jianjun Sun

## Abstract

*Mycobacterium tuberculosis* virulence factors, EsxA and EsxB, are secreted as a heterodimer (EsxA:B) and play an essential role in mycobacterial phagosomal escape and cytosolic translocation. Current studies support a model that EsxA must dissociate from its chaperon EsxB at low pH in order for EsxA to interact with host membranes. However, the mechanism of the heterodimer separation is not clear. In the present study, we have obtained evidence that the N^α^-acetylation of Thr2 on EsxA, a post-translational modification that is present in mycobacteria, but absent in *E. coli*, is required for the heterodimer separation. The point mutations at Thr2 without N^α^-acetylation inhibited the heterodimer separation and hence prevented EsxA from interacting with the membranes, which resulted in attenuated mycobacterial cytosolic translocation and virulence. Molecular dynamic simulation showed that at low pH the N^α^-acetylated Thr2 made direct and frequent “bind-and-release” contacts with EsxB, which generates a dragging force to pull EsxB away from EsxA.

## Introduction

*Mycobacterium tuberculosis* (Mtb) is the causative agent for tuberculosis (TB), one of the leading infectious diseases in the world with 10 million people falling ill in 2017 and approximately 1.6 million deaths (1, 2). It is believed that after the Mtb-containing aerosolized droplets are inhaled into the lung, Mtb is encountered by alveolar macrophages and internalized into the phagosome, where Mtb manages to survive through arresting phagosome maturation, including inhibition of vATPase-mediated acidification (3–6). Recent compelling evidence support that Mtb penetrates the phagosome and translocate into the cytosol (termed cytosolic translocation), where Mtb replicates and undergoes cell-to-cell spreading (7). The ability of Mtb to arrest phagosome maturation and to translocate from the phagosome to the cytosol has been attributed, at least in part, to the Type VII secretion system, named ESX-1 and the secreted virulence factors EsxA (ESAT-6) and EsxB (CFP-10). The Mtb mutants with either gene deletions or defects in secretion of EsxA and/or EsxB, were not able to translocate into the cytosol and showed significant reduction in host cell lysis and cell-to-cell spreading (7–19).

In our previous studies, we have found that Mtb EsxA exhibits a unique membrane-permeabilizing activity that is not present in the homologous EsxA ortholog from non-pathogenic *Mycobacterium smegmatis* (Ms) (20). EsxA undergoes pH-dependent conformational changes, inserts into the membrane and forms a membrane-spanning complex (21). The essential role of EsxA membrane-permeabilizing activity in phagosome rupture and cytosolic translocation is further confirmed by a recent study, in which single-residue mutations at Gln5 (Q5) of EsxA up or down regulated the membrane-permeabilizing activity and consequently up or down regulated mycobacterial cytosolic translocation and virulence in cultured cells and in zebra fish (22).

The genes encoding EsxA and EsxB are located in the same operon within the ESX-1 locus. EsxA and EsxB are co-expressed and co-secreted as a heterodimer (23). Our earlier study has demonstrated that EsxA, but not EsxB, has the membrane-permeabilizing activity, and EsxB is believed to function as a chaperon (20). Current studies support a model that the heterodimer is dissociated at low pH to allow EsxA to penetrate the membranes (24). However, the data regarding to the heterodimer dissociation are conflicting. The native heterodimer extracted from Mtb culture filtrate was found to be dissociated at low pH (24). Surprisingly, however, the studies using the recombinant proteins prepared from *E. coli* suggest that the heterodimer was not dissociated by acidification. This is evidenced by one of our earlier studies that the heterodimer prepared from *E. coli* (now termed Ec-heterodimer) was inactive in membrane disruption. In the absence of lipid membranes, EsxA formed aggregates in the acidic solution due to increased solvent-exposed hydrophobicity. In contrast, the Ec-heterodimer showed little aggregation at pH 4.0, suggesting that EsxB remains bound to EsxA at low pH and prevents EsxA from forming aggregates, which otherwise would be observed if EsxA was released from EsxB (20). Our data are consistent with an earlier CD analysis showing that the Ec-heterodimer is not dissociated at low pH (25).

We hypothesized that the mycobacteria-produced proteins contain unique features (e.g. post-translational modifications, PTMs) that are required for heterodimer dissociation at low pH. In line with this hypothesis, the native EsxA protein isolated from the culture filtrate of Mtb was displayed as multiple spots in 2D SDS-PAGE, and some of the spots contained a N^α^-acetylation at the residue Thr2 (26). Moreover, the heterodimer produced from a Ms strain was found to have a N^α^-acetylation on the Thr2 residue of EsxA (27). Interestingly, EsxB preferred to bind the non-acetylated EsxA, but not the acetylated form in a 2-D overlay assay (26). Deletion of the N^α^-acetyltransferase in *Mycobacterium marinum* (Mm) disrupted the homeostasis of EsxA N^α^-acetylation and attenuated the virulence (28). Together, these studies suggest that the N^α^-acetylation of EsxA plays an important role in mycobacterial virulence through facilitating heterodimer dissociation at low pH.

In the present study, we have obtained the evidence showing that the N^α^-acetylation at Thr2 of EsxA is required for EsxA membrane permeabilization, mycobacterial cytosolic translocation and virulence through facilitating heterodimer dissociation.

## Experimental Procedures

### Generation of T2X mutations on the *esxA* gene for expression in *E. coli*, *M. smegmatis* and *M. marinum*

#### For expression in *E. coli*

Using the previously reported plasmid pET22b-esxA-His_6_ as the template (20–22, 29), the mutations T2A, T2Q, T2R, and T2S were introduced into the *esxA* gene by PCR using the primers listed in **Table S1**. All of the mutations were confirmed by DNA sequencing. The resultant plasmids were transformed into BL21 (DE3) cells for expression. The cells were grown at 37 °C while shaking at 250 rpm until OD_600_ reached 0.6-0.8. Protein expression was induced by adding 1 mM isopropyl-β-D-1-thiogalactopyranoside (IPTG) for 3-8 h at 37 °C. The cells were harvested and the proteins were purified as previously described (20–22, 29).

#### For expression in Ms

The pMyNT plasmid containing the *esxB-esxA* operon (27) was used as the template. The mutations T2A, T2Q, T2R and T2S were introduced by overlapping PCR using the primers listed in **Table S1**. All of the mutations were confirmed by DNA sequencing. The pMyNT plasmids carrying various T2X mutations were electroporated into Ms mc^2^155 strain (voltage: 2,500 V, capacitance: 25 μF, resistance: 1,000 Ω). The Ms cultures were grown at 37 °C overnight or until OD_600_ reached 2.0. Protein expression was induced by adding 0.2% (w/v) acetamide for 12-16 h. The cells were harvested. The soluble heterodimer proteins were purified with immobilized metal ion affinity chromatography by passing through a Ni^2+^-column, followed by a sizing exclusion chromatography as previously described (27, 30–33).

#### For expression in Mm

The T2X mutations (T2A, T2R, T2Q and T2S) were generated by site-directed mutagenesis (Agilent Quick Change Kit) using the pMH406 plasmid containing *esxB-esxA* operon as a template. The mutations were confirmed by DNA sequencing. The mutated plasmids were electroporated into MmΔEsxA:B as previously described (32).

### Liposome leakage assay

The liposome leakage assay was performed as previously described (20, 21, 29, 34, 35). Briefly, 20 mg of DOPC (1,2-dioleoyl-sn-glycero-3-phosphocholine) was dried with nitrogen air and left in a vacuum overnight. The samples were rehydrated with 1 ml of the buffer (5 mM HEPES, 50 mM ANTS (8-aminonapthalene-1,3,6 trisulfonic acid) and 50 mM DPX (p-xylene-bispyridinium bromide)). The suspension was subjected to 6x freeze-thaw cycles and extruded via a 0.2 μm membrane filter for 20 times. The liposomes were then desalted to remove excess ANTS and DPX using a Hi-trap desalting column. The desalted liposomes were mixed with 150 mM NaCl, 100 mM NaAc at pH 4.0. The reaction mixture was excited at 350 nm and emissions were recorded at 520 nm in an ISS K2 phase modulation fluorometer. 100 μg of the tested protein was injected into the solution after approximately 30 s of the assay and the fluorescence was recorded.

### Isolation of EsxA proteins from the Ms-produced EsxA:B heterodimer

The EsxA:B heterodimer purified from Ms was incubated in a solution containing 6 M guanidine at 4 μC overnight. The proteins were then passed through a HisTrap column (GE Healthcare). The His-tagged EsxB protein was bound to the column, and the un-tagged EsxA was collected in flow through. The His-tagged EsxB protein was eluted by an imidazole gradient. Both EsxA and EsxB were subjected to an extensive dialysis using a 3,000 MWCO membrane. The samples were then concentrated and passed through gel filtration for a complete buffer exchange.

### Detection of N^α^-acetylation by NBD-Cl

NBD–Cl (4-chloro-7-nitrobenzof-urazan) only reacts with free N-terminal α-amino group in non-acetylated proteins and emits fluorescence, and it does not react with N^α^-acetylated proteins due to lack of free N-terminal amino group (36). Proteins (6 μM) were incubated with 0.5 mM NBD-Cl at room temperature. At different hours of post-incubation, the samples were subjected to fluorescence measurement with excitation at 460 nm and emission at 535 nm.

### Identification of post-translational modifications by mass spectrometry

The purified proteins were digested by FASP-trypsin and analyzed by LC-MS/MS on a QE-Classic. The results were analyzed by PD2.1 search against a combined database containing *E. coli* BL21 and Ms mc^2^155.

### Western Blotting

The Mm strains were cultured in 7H9 medium and grown to mid-log phase. They were washed with PBS and transferred to Sauton’s medium while normalizing all cultures to OD_600_ = 0.8. The bacteria were cultured for 2 days until harvest. The bacterial cells were collected by centrifugation. The proteins in the culture supernatant were precipitated by trichloroacetic acid (TCA). The bacterial cells were resuspended in 1 ml of PBS containing a cocktail of protease inhibitors (Thermofisher) and sonicated at 30% amplitude for 5 cycles of 30 s pulse and 60 s rest. The culture filtrates and total bacterial lysates were applied to SDS-PAGE and transferred onto PVDF membrane. Western blots were performed to detect EsxA using anti-EsxA antibody (sc-57730, Santa Cruz). As controls, Ag85 (secreted in culture filtrate) and GroEL (only in cell lysate) were also detected by anti-Ag85 (NR-13800, BEI) and anti-GroEl antibodies (NR-13813, BEI), respectively.

### Live/Dead Cytotoxicity Assay

RAW264.7 cells were cultured in DMEM medium containing 10% fetal bovine serum (FBS) with penicillin and streptomycin (100U/mL) at 37 °C and 5% CO_2_. Raw 264.7 macrophages were plated in a 24-well plate with a density of 5 ×10^5^/well for infection on the following day. The Mm strains were prepared with a single cell preparation protocol as previously described (22, 37). RAW264.7 cells were infected with various Mm strains at a multiplicity of infection (MOI) of 10 for 1 h. The macrophages were washed 3 times with PBS to remove free mycobacteria and incubated for another 3 h. The macrophages were stained using Calcein-AM and ethidium homodimer (Life Sciences) for 30 min, enabling visualization under a fluorescence microscope for green cells (live) and red cells (dead). The numbers of dead cells were quantified from dozens of random fields from each sample.

### CCF-4 FRET assay

Mycobacterial cytosolic translocation was measured by CCF-4 FRET assay as previously described (17, 22, 38). Briefly, RAW264.7 cells were plated in a 6-well plate at a density of 2.5 × 10^6^ cells/well. The macrophages were then incubated with CCF4-AM according to the manufacturer’s protocol (Liveblazer B/G loading kit, Life Sciences). Cells were infected at a MOI of 10 for 2 h. Following infection, macrophages were washed 3 times using PBS. DMEM media with 10% FBS was added to the cells and incubated for approximately 2 days. The samples were then excited at 409 nm and the emissions were measured at 450 nm and 535 nm. The blue/green ratio was calculated as I_450_/I_535_.

### Molecular dynamic simulation

The structure of EsxA:B heterodimer was downloaded from Protein Data Bank with PBD ID 1WA8 (39). DelPhiPka web server (40) was used to obtain the protonation states of ionizable residues at pH 4 and pH 7 and assign the respective states with Visual Molecular Dynamics (VMD) (41). N^α^-acetylation of Thr2 was performed on VMD after removal of Met1. The four structures i.e. non-acetylated and acetylated of EsxA:B heterodimer at either pH 7 or pH 4 were then solvated in water box with TIP3 (42) water model and ionized with 150 mM NaCl in VMD. The final systems were then simulated with molecular dynamics (MD) simulation program NAMD (43). Each simulation was performed for 20 ns employing force field CHARMM27 (44). The temperature was set as 300 K and the pressure was 1 atm. The snapshots from the simulations were taken to study the behaviors of the N-terminal loop of EsxA with and without N^α^-acetylation of Thr2 at pH 4 and pH 7.

## Results

### The Ms-produced heterodimer, but not Ec-produced heterodimer, disrupted liposomal membrane at low pH

We hypothesized that Ms-heterodimer, but not Ec-heterodimer, dissociates at low pH to allow EsxA permeabilize the liposomal membrane. The membrane-permeabilizing activity of Ms-heterodimer and Ec-heterodimer was tested with the ANTS/DPX fluorescence de-quenching assay. As expected, Ms-heterodimer permeabilized the membrane at low pH, while Ec-heterodimer did not (**Fig. 1A**). NBD–Cl (4-chloro-7-nitrobenzof-urazan) only reacts with free N-terminal α-amino group in non-acetylated proteins and emits fluorescence, but it does not react with N^α^-acetylated proteins due to lack of free N-terminal amino group. Thus, we used NBD-Cl to test the states of N^α^-acetylation for Ms-heterodimer and Ec-heterodimer. As expected, the Ms-heterodimer exhibited a significantly lower NBD-Cl fluorescence, compared to Ec-heterodimer, indicating that Ms-heterodimer, but not Ec-heterodimer, is N^α^-acetylated (**Fig. 1B**).

**Figure 1.**
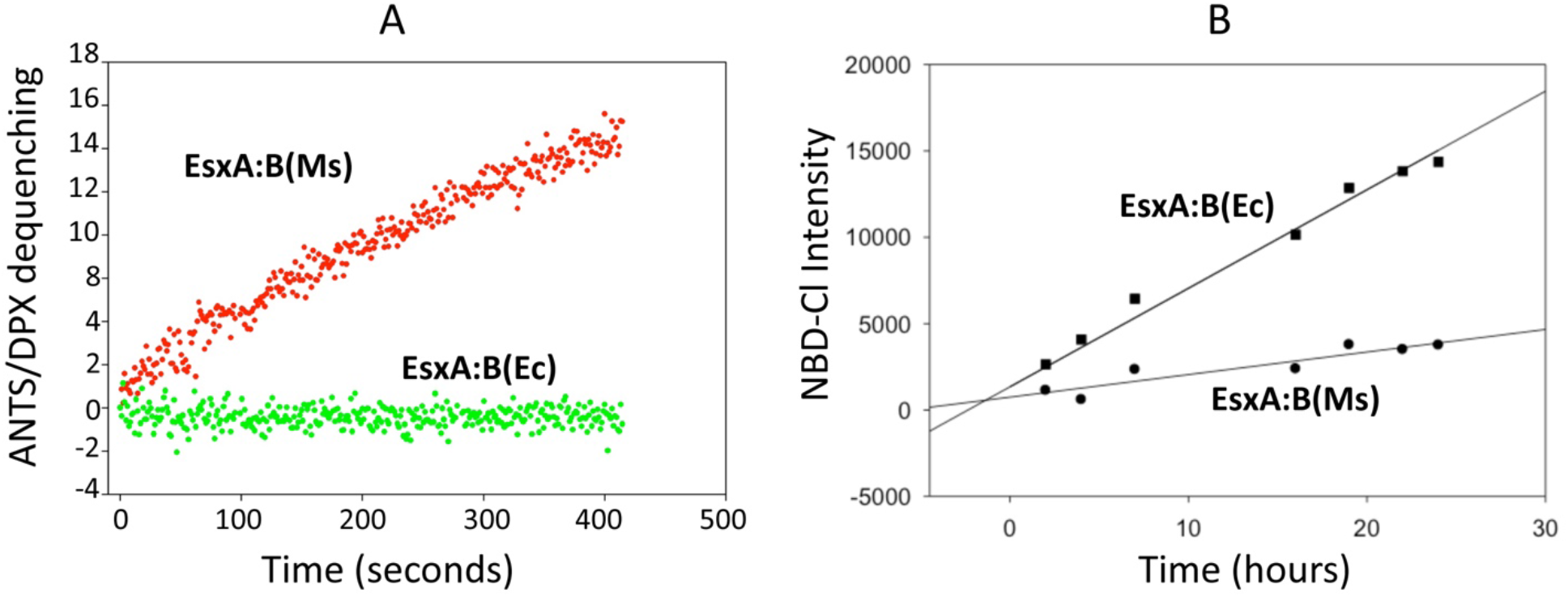
The EsxA:B heterodimer produced in Ms, but not in *E. coli*, permeabilized the liposomes at low pH, implicating the role of N^α^-acetylation in heterodimer dissociation. **A.** The heterodimer EsxA:B purified from Ms or *E. coli* were tested with ANTS/DPX dequenching assay at pH 4.0. **B.** Equal amount of the heterodimer EsxA:B(Ms) or EsxA:B(Ec) were incubated with NBD-Cl at room temperature. NBD-Cl is a fluorescent dye that only react with a free N-terminal site without N^α^-acetylation was recorded at 535 nm (excitation at 480 nm) and plotted as a function of time.

### The mutations at Thr2 abolished the membrane-permeabilizing activity of the Ms-heterodimers through blocking separation of EsxA and EsxB

The Q and A residues have been used to functionally mimic acetylation of amino group of an internal K residue, while R serves as a non-acetylated control (45). Thus, we generated T2A, T2Q and T2R mutations and tested the effects of these mutations on the heterodimer membrane-permeabilizing activity.

Unexpectedly, all of the mutations abolished the Ms-heterodimer’s membrane-permeabilizing activity (**Fig. 2A and B**). The result suggests that either the mutations blocked the heterodimer separation or abolished the EsxA membrane-permeabilizing activity. To exclude the possibility that the mutations abolished the membrane-permeabilizing activity, we purified the EsxA proteins containing the same mutations from *E. coli* and applied them to ANTS/DPX dequenching assay. The result showed that the mutations did not affect EsxA membrane-permeabilizing activity (**Fig. 2C and D**), suggesting that the mutations at Thr2 blocked the heterodimer separation at low pH.

**Figure 2.**
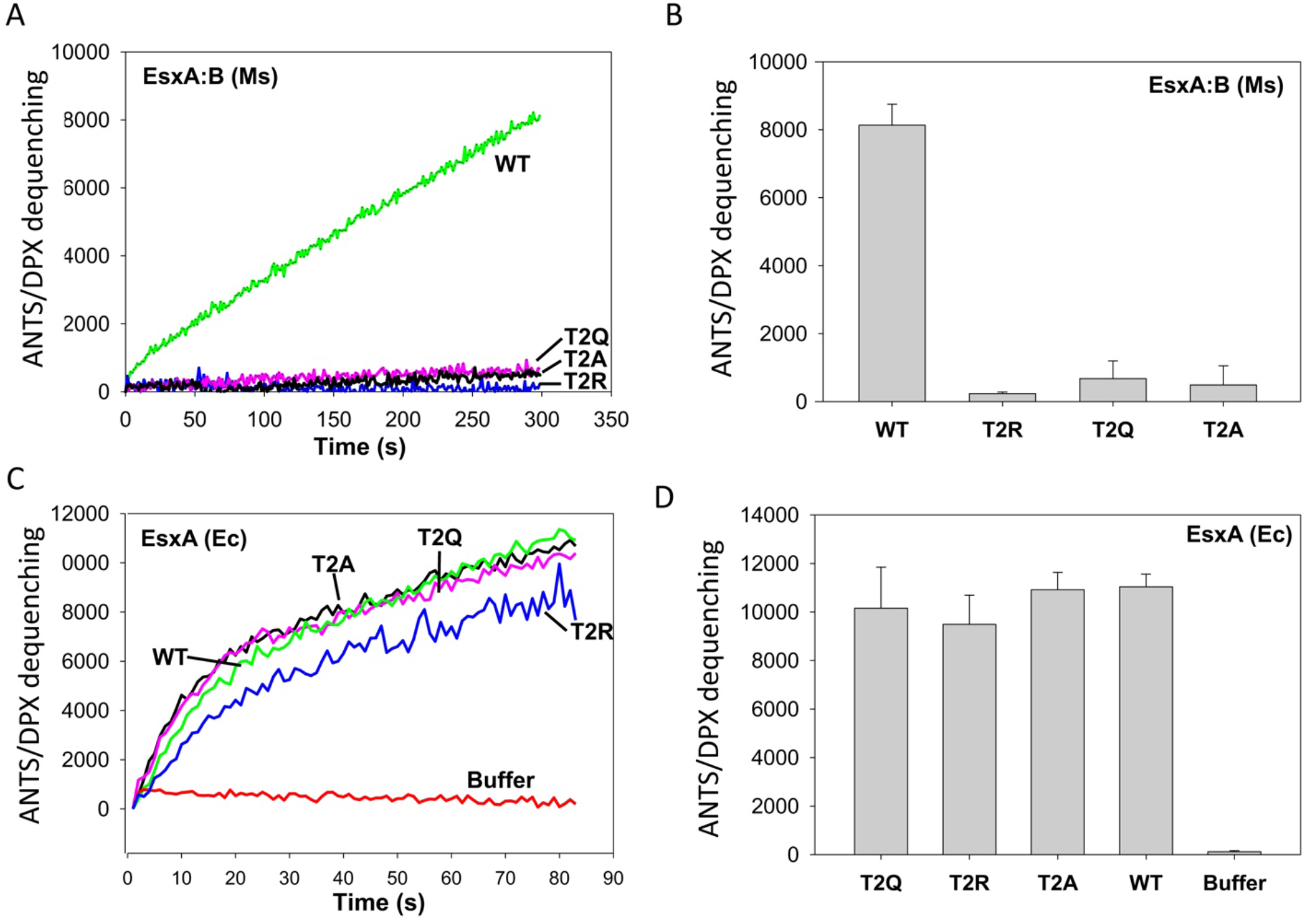
The mutations at Thr2 of EsxA diminished the membrane-permeabilizing activity of EsxA:B heterodimer. **A.** The EsxA:B heterodimer proteins (wild type and the mutants carrying mutations T2A, T2Q and T2R) were purified from Ms. The membrane-permeabilizing activity of the purified heterodimer proteins was tested by ANTS/DPX fluorescence dequenching assay. The representative ANTS/DPX fluorescent dequenching curves were shown. **B.** The average end-point fluorescence intensities from at least three independent experiments were calculated. **C.** The EsxA proteins (WT and T2A, T2Q and T2R mutants) were purified from Ec. The membrane-permeabilizing activity of the Ec-EsxA proteins purified from Ec was tested by ANTS/PDX assay. The representative curves were shown. **D.** The average end-point fluorescence intensities from at least 3 independent experiments were calculated and shown.

### The mutations that block separation of EsxA and EsxB do not have N^α^-acetylation

To validate the acetylation state of EsxA wild type and the mutants, we developed a protocol and isolated EsxA and EsxB from the Ms-heterodimers (**Fig. 3A**). The presence of N^α^-acetylation on the EsxA proteins was measured by NBD-Cl (**Fig. 3B**). Consistent with the results in **Figure 1B**, Ec-EsxA(WT) had a significantly higher fluorescence signal than Ms-EsxA(WT). Similar to Ec-EsxA(WT), the Ms-EsxA mutants (T2A, T2Q, and T2R) emitted significantly higher fluorescence signals than Ms-EsxA(WT), suggesting that the Ms-EsxA mutants were not N^α^-acetylated. Next, we applied Ms-EsxA(WT), Ms-EsxA(T2A) and Ec-EsxA(WT) to LC-MS/MS to further confirm the N^α^-acetylation states. In addition, Ms-EsxA(T2S), a mutant equivalent to WT, was included in the LC-MS/MS analysis. The results showed that both Ms-EsxA(WT) and Ms-EsxA(T2S) had the first Met residue removed and the second residue (either Thr2 or Ser2) acetylated (**Fig. 3C**). While Ms-EsxA(T2A) had the first Met residue removed, the second Ala residue was not acetylated. The Ec-EsxA(WT) still has the first Met residue and no acetylation on Thr2. Interestingly, the LC-MS/MS detected multiple acetylation and oxygenation modifications in the internal sequences of both Ms-EsxA and Ec-EsxA and the roles of these modifications are currently unknown.

**Figure 3.**
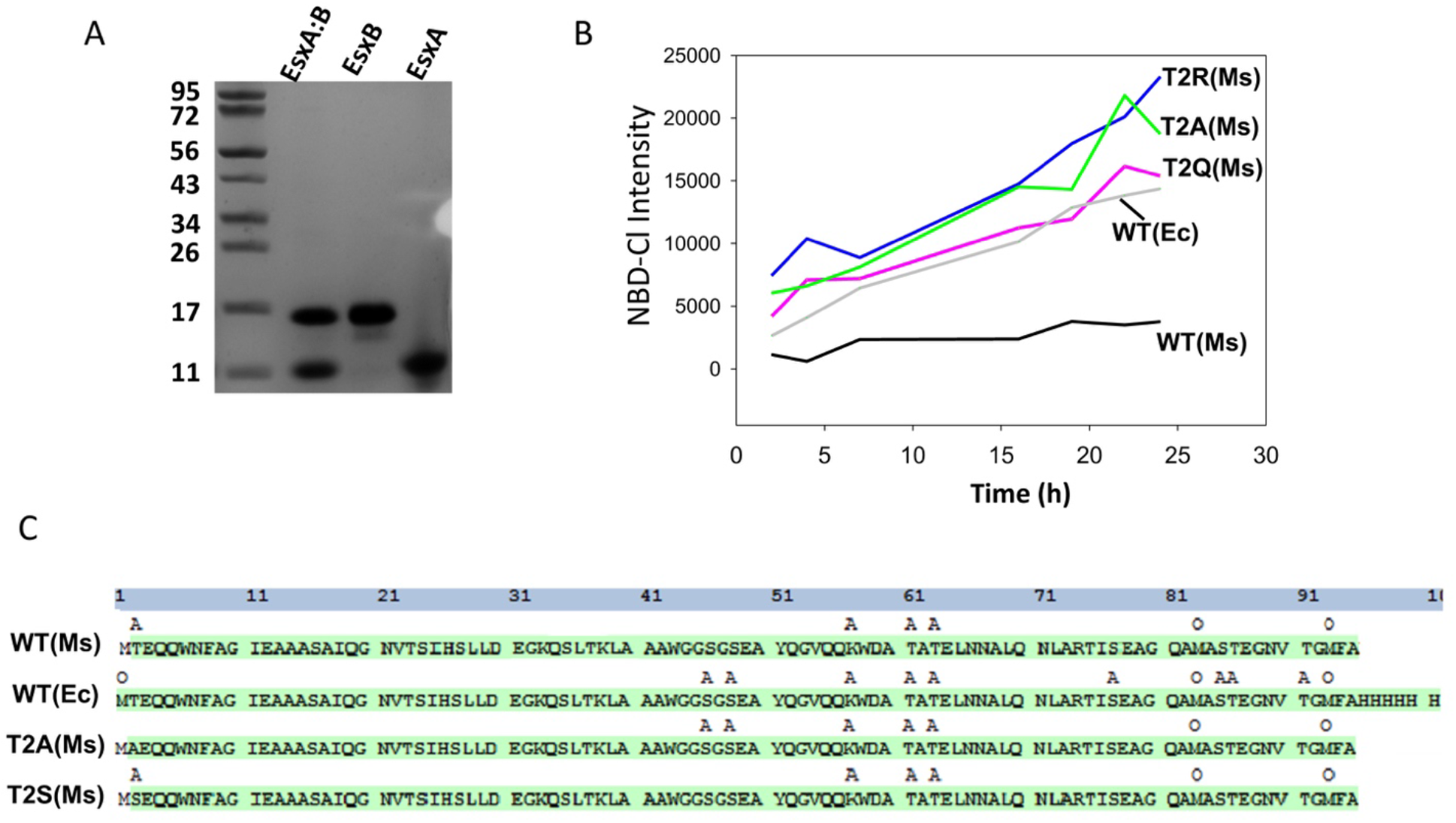
Detection of N^α^-acetylation of EsxA by NBD-Cl and mass spectrometry. **A.** The EsxA:B heterodimer purified from *M. smegmatis* was separated by 6 M guanidine, after which EsxA and EsxB were purified separately as described in Experimental Procedures. **B**. The indicated EsxA proteins were incubated with NBD-Cl. At the indicated times, the fluorescence intensity of NBD-Cl was measured. **C.** The indicated EsxA proteins were analyzed via LC/MS/MS to identify the post-translational modifications. The residues with acetylation were labeled as “A” and the residues with oxidation were labeled as “O”.

### EsxB preferred to bind the non-acetylated EsxA to inhibit the membrane-permeabilizing activity

An earlier study has shown that EsxB preferred to bind the non-acetylated EsxA than acetylated EsxA in a 2D overlay assay (26). Thus, we hypothesize that EsxB will prefer to inhibit the membrane-permeabilizing activity of the non-acetylated Ms-EsxA(T2A) than that of the acetylated Ms-EsxA(WT). First, we tested and confirmed that the proteins Ms-EsxA(WT) and Ms-EsxA(T2A) that were isolated from the heterodimers had similar membrane-permeabilizing activity to Ec-EsxA(WT), which once again confirms that the states of N^α^-acetylation does not affect membrane-permeabilizing activity (**Fig. 4A and B**). Then, Ms-EsxA(WT) and Ms-EsxA(T2B) were incubated with EsxB at a series of EsxB/EsxA molar ratios. As expected, EsxB preferred to inhibit Ms-EsxA(T2A) than Ms-EsxA(WT) in membrane permeabilization, especially at the lower EsxB/EsxA ratios (**Fig. 5A, B and C**).

**Figure 4.**
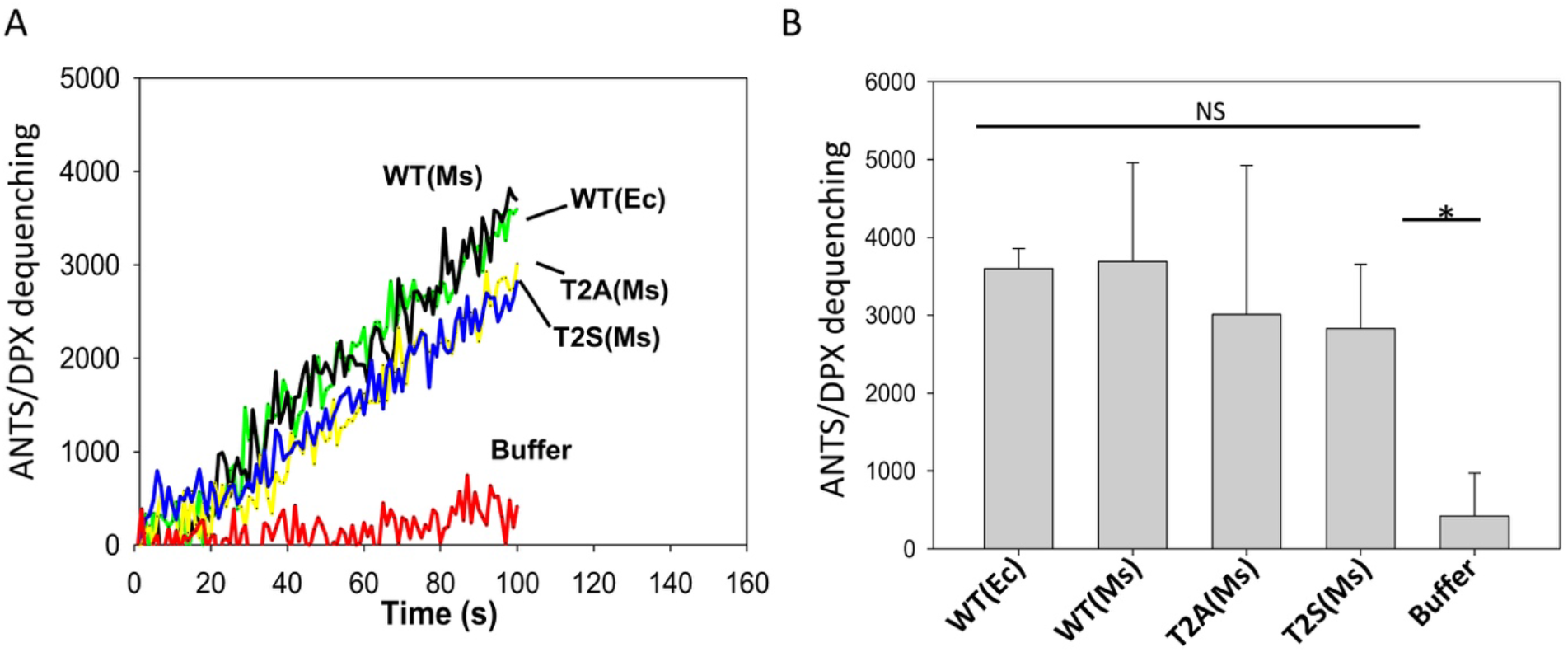
The N^α^-acetylation of EsxA did not affect the membrane-permeabilizing activity of EsxA. **A.** The indicated Ms-EsxA proteins isolated from the Ms-heterodimer were tested for membrane-permeabilizing activity using ANTS/DPX assay. The Ec-EsxA (WT) protein was used as a control. The representative curves from at least three independent experiments were shown. **B.** The average end-point fluorescence intensity from at least three independent experiments was calculated and shown.

**Figure 5.**
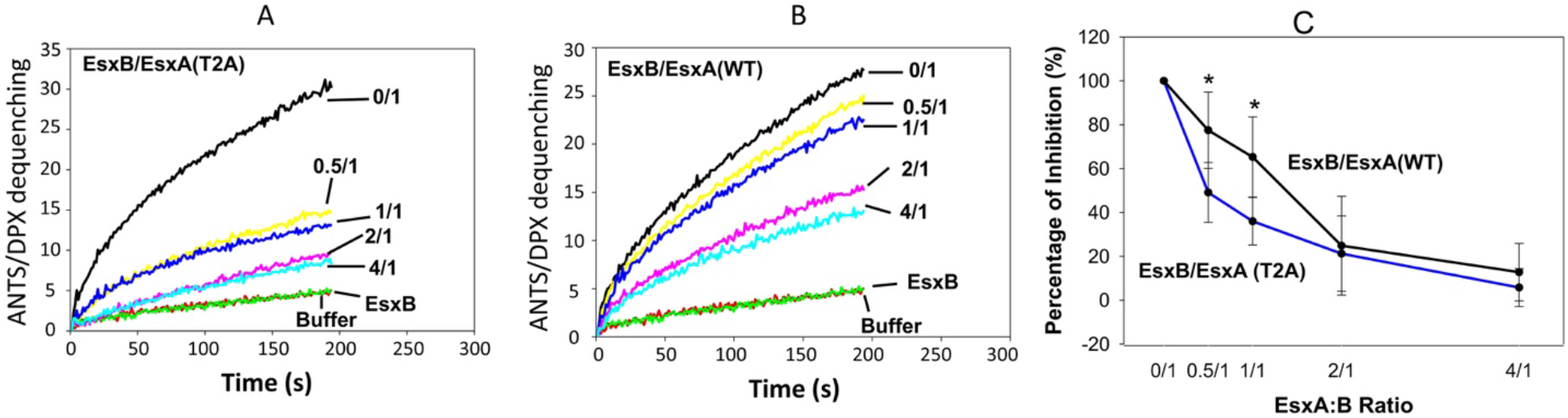
EsxB preferentially inhibited non-acetylated EsxA(T2A) over its N^α^-acetylated counterpart. The Ms-produced EsxA(WT) and EsxA(T2A) proteins were incubated with various concentrations of EsxB as the indicated molar ratios. The mixtures were tested by ANTS/DPX assay for membrane-permeabilizing activity. The representative curves were shown in **A** and **B**, respectively. The relative inhibition from at least three independent experiments was summarized in **C**.

### The mutations without N^α^-acetylation attenuated mycobacterial virulence and inhibited cytosolic translocation

Here, we investigated the effects of the T2X mutations in mycobacterial pathogenesis. The genes carrying T2X mutations were expressed in the MmΔEsxA:B strain, in which the endogenous *esxB-esxA* operon was deleted. We found that the T2X mutations did not affect the expression and secretion of EsxA and EsxB in the Mm strains (**Fig. 6A**). As expected, the Mm strains carrying the non-acetylated mutations T2A, T2Q and T2R had a significantly lower cytotoxicity than the strain carrying the acetylated mutation T2S (**Fig. 6B**). MmWT and MmΔEsxA:B were used as the positive control and negative control, respectively. Furthermore, using the previously established CCF4-FRET assay, we found that the non-acetylated mutations T2A, T2Q and T2R abolished mycobacterial cytosolic translocation, while T2S maintained a similar activity as the wild type (**Fig. 6C**). The data is consistent to the previous report that deletion of a N^α^-acetyltransferase in Mm disrupted the homeostasis of EsxA acetylation and attenuated the virulence (28).

**Figure 6.**
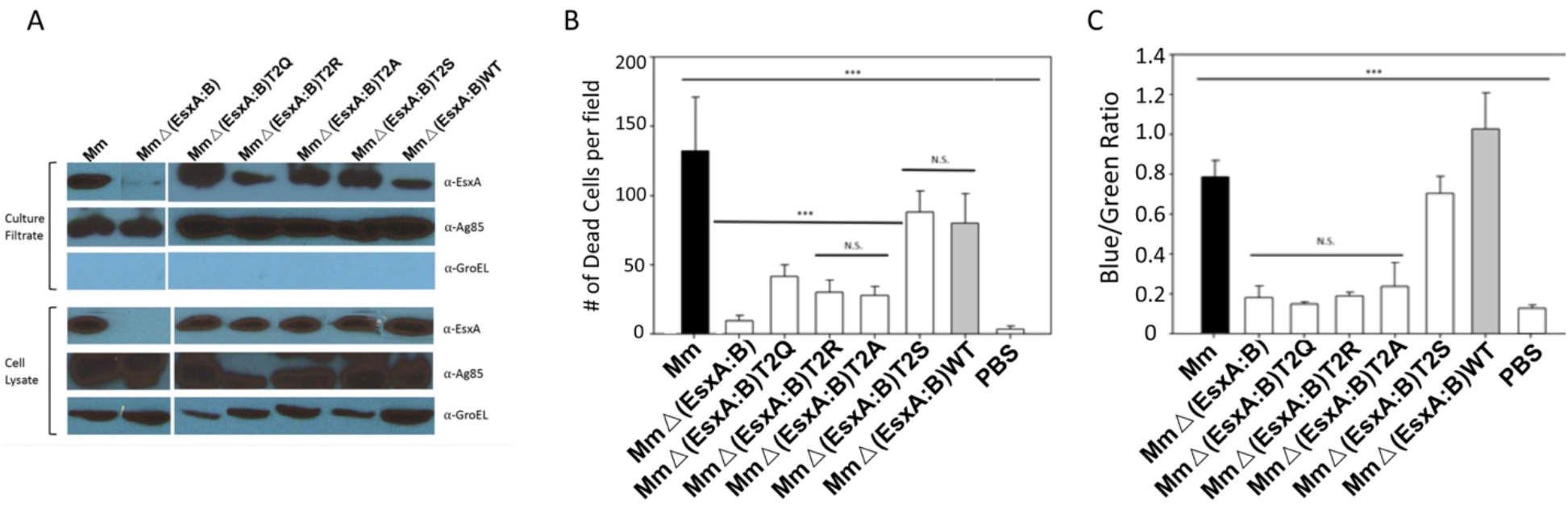
The non-N^α^-acetylate EsxA diminished mycobacterial virulence and cytosolic translocation in macrophages. **A.** The EsxA proteins in the total cell lysates or in the culture filtrate of the indicated Mm strains were detected by western blots using anti-EsxA antibody. GroEL was detected by anti-GroEL antibody as a loading control for cell lysate. Ag85 was detected by anti-Ag85 as a loading control for culture filtrate. **B.** RAW263.4 cells were infected with the indicated Mm strains at MOI of 10. The cytotoxicity was measured by using the Live/Dead assay. Dead cells were counted in random fields (***: n =22, p <0.05). **C**. Mycobacterial cytosolic translocation was monitored by using CCF4-AM as a FRET reporter. The Blue/Green ratio was measured by comparing emissions at 450nm and 530nm with excitation at 409 nm. The data were calculated from at least three independent experiments (***: p <0.05).

### Molecular dynamic simulation detects frequent “bind-and-release” contacts between the acetylated Thr2(Ac) and EsxB

The reported NMR solution structure of EsxA:B heterodimer does not have the N^α^-acetylation on Thr2, and the structure shows Thr2 is distal from the contact interface between EsxA and EsxB. It is not clear how the acetylation at Thr2 affects the heterodimer separation at low pH. Thus, we performed molecular dynamic (MD) simulation on the heterodimers containing either non-acetylated or acetylated Thr2 at pH 7 and pH 4, respectively (**Figure 7**). At pH 7 the non-acetylated Thr2 comes in a close vicinity of EsxB, but it is unable to make a direct contact to EsxB (**Figure 7A, C**). Compared to non-acetylated Thr2, the acetylated Thr2(Ac) moves further away from EsxB at pH 7 (**Figure 7B, D**). Interestingly, at pH 4 the non-acetylated N-terminal loop of EsxA has no direct contact with EsxB (**Figure 7E, G**), but the acetylated N-terminal loop is able to make direct contacts with EsxB (**Figure 7F, H**). The MD simulation shows that the acetylated loop make direct contacts with EsxB in a frequent “bind-and-release” mode **(Movie S1)**), which generates a pulling force to trigger the dissociation of the complex.

**Figure 7.**
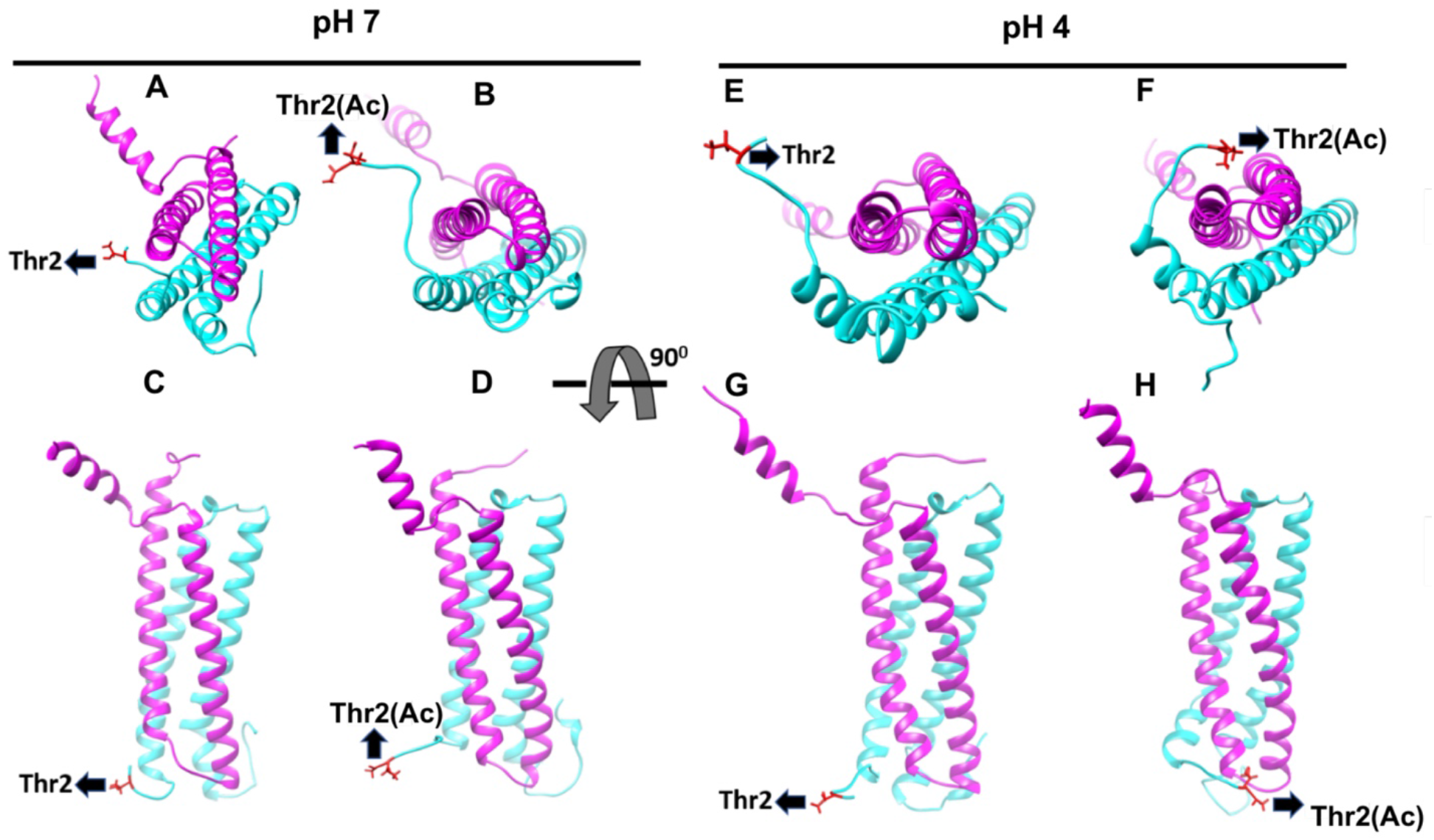
Molecular dynamic simulation detects the acetylated Thr2(Ac) interacts with EsxB in a “bind-and-release” mode. The structures of EsxA:B heterodimers with/out N^α^-acetylation were analyzed by molecular dynamic simulation. The figures were generated from snapshots of 20 ns MD simulations at pH 7 and pH 4. EsxA, EsxB and the Thr2 residue are shown in cyan, pink and red, respectively. The structures of EsxA:B heterodimer with non-acetylated Thr2 at pH 7 and pH 4 are shown in **A** (pH 7, top view), **C** pH 7, side view), **E** (pH 4, top view) and **G** (pH 4, side view), respectively. The structures of EsxA:B heterodimer with acetylated Thr2(Ac) are shown in **B** (pH 7, top view), **D** (pH 7, side view), **F** (pH 4, top view) and **H** (pH 4, side view), respectively.

## Discussion

EsxA and EsxB are co-expressed and co-secreted as a heterodimer in mycobacteria. The role of ESX-1, EsxA and EsxB in mycobacterial cytosolic translocation and virulence has been confirmed in a series of studies (7, 17–19). Earlier biochemical studies have demonstrated that EsxA has a pH-dependent membrane-permeabilizing activity, while EsxB appears to function as a chaperone for EsxA (20). Current studies support a model that the EsxA:B heterodimer is dissociated at low pH, which allows EsxA to permeabilize the membranes (24). However, the mechanism of the heterodimer separation is not known. For the first time, the present study tested the heterodimers with or without N^α^-acetylation of Thr2 and obtained the evidence that the N^α^-acetylation at Thr2 of EsxA facilitates the heterodimer separation at low pH, which allows EsxA to permeabilize liposomal membrane in vitro as well as mediate mycobacterial phagosome escape and cytosolic translocation in mycobacteria-infected macrophages.

As discussed above, the essentiality of EsxA and EsxB in mycobacterial pathogenesis has been well documented in a series of reports. Genetic manipulations that either deleted the gene of *esxA* or *esxB*, or abolished the secretion of EsxA and EsxB, have attenuated mycobacterial virulence and inhibited the phagosome rupture, cytosolic translocation and cell-to-cell spreading (7, 10, 17–19). Moreover, the biochemical characterizations have demonstrated that EsxA possesses a unique membrane-permeabilizing activity that is not present in its ortholog in non-pathogenic *M. smegmatis* (20). Thus, it is reasonable to believe that during the course of infection, the secreted EsxA exerts its membrane-permeabilizing activity to penetrate the phagosome membranes and facilitate mycobacterial cytosolic translocation. We have reported that the mutations at the Gln 5 residue of EsxA (e.g. Q5V and Q5K) have resulted in up- or down-regulation of EsxA membrane-permeabilizing activity *in vitro*. Moreover, these mutations up- or down-regulated the mycobacterial virulence and cytosolic translocation accordingly, demonstrating the specific and accurate correlation between EsxA membrane-permeabilizing activity and mycobacterial virulence as well as the ability to penetrate phagosome membrane (22). Once again, the present study provides new evidence that the N^α^-acetylation at Thr2 of EsxA is required for mycobacterial virulence and cytosolic translocation through facilitating the heterodimer separation.

Since Thr2 has no direct contact with EsxB as shown in the reported solution structure of EsxA:B heterodimer, how N^α^-acetylation on Thr2 affect heterodimer separation had become a puzzle. Here, the MD simulation result has provided a convincing evidence that the acetylated Thr2(Ac) makes frequent “bind-and-release” contacts with EsxB only at low pH, generating a dragging force to pull EsxB away from EsxA.

A recent study has shown that the recombinant EsxA does not lyse cell membranes, and the lytic activity previously attributed to EsxA is due to residual ASB-14 detergent in the preparation (46). In fact, we had the similar observations that addition of the recombinant EsxA protein to the surface of lung epithelial cell lines WI-26 and A549 did not lyse the cells (data not shown). Moreover, Conrad et. al. showed that blocking phagosomal acidification by Bafilomycin did not decrease the ESX-1-mediated phagosomal permeabilization, suggesting that acidification is not required for membrane permeabilization (46). It is not clear how the discrepancy arises and what is the broken link between the EsxA pH-dependent membrane-permeabilizing activity in model membrane and the ability of mycobacteria to rupture phagosome membrane during infection. Other factors from mycobacteria and host cells, even including properties of target membranes, may be involved in this process, which warrants further investigations.

## Acknowledgement

We thank Dr. Matthias Wilmanns for providing the pMyNT plasmid. The following reagents were obtained through BEI Resources, NIAID, NIH: Polyclonal Anti-*Mycobacterium tuberculosis* Antigen 85 Complex (FbpA/FbpB/FbpC; Genes Rv3804c, Rv1886c, Rv0129c) (antiserum, Rabbit), NR-13800; Monoclonal Anti-*Mycobacterium tuberculosis* GroEL2 (Gene Rv0440), Clone CS-44 (produced *in vitro*), NR-13813.

The study is supported by the grants from NIGMS (SC1GM095475 to J. Sun), National Center for Research Resources (5G12RR008124) and National Institute on Minority Health and Health Disparities (G12MD007592). The content is solely the responsibilty of the authors and does not necessarily represent the official views of the National Institutes of Health.

